# Inositol pyrophosphate-controlled kinetochore architecture and mitotic entry in *S. pombe*

**DOI:** 10.1101/2022.08.08.503146

**Authors:** Natascha Andrea Kuenzel, Abel R. Alcázar-Román, Adolfo Saiardi, Simon M. Bartsch, Sarune Daunaraviciute, Dorothea Fiedler, Ursula Fleig

## Abstract

Inositol pyrophosphates (IPPs) comprise a specific class of signaling molecules that regulate central biological processes in eukaryotes. The conserved Vip1/PPIP5K family controls intracellular IP_8_ levels, the highest phosphorylated form of IPPs present in yeasts, as it has both inositol kinase and pyrophosphatase activities. Previous studies have shown that the fission yeast *S. pombe* Vip1/PPIP5K family member Asp1 impacts chromosome transmission fidelity via modulation of spindle function. We now demonstrate that an IP_8_ analogue is targeted by endogenous Asp1 and that cellular IP_8_ is subject to cell cycle control. Mitotic entry requires Asp1 kinase function and IP_8_ levels are increased at the G2/M transition. In addition, the kinetochore, the conductor of chromosome segregation assembled on chromosomes is modulated by IP_8_. Members of the yeast CCAN kinetochore-subcomplex such as Mal2/CENP-O localize to the kinetochore depending on the intracellular IP_8_-level: higher than wild-type IP_8_ levels reduces Mal2 kinetochore targeting, while a reduction in IP_8_ has the opposite effect. As our perturbations of the inositol polyphosphate and IPP pathways demonstrate that kinetochore architecture depends solely on IP_8_ and not on other IPPs, we conclude that chromosome transmission fidelity is controlled by IP_8_ via an interplay between entry into mitosis, kinetochore architecture and spindle dynamics.

## Introduction

Inositol pyrophosphates (IPPs) are highly energetic molecules derived from *myo*-inositol carrying monophosphates and one or two diphosphate groups at defined positions of the inositol ring. IPPs are present in all eukaryotes and were categorized as signaling molecules due to their rapid turnover [1]. The functions of these molecules are very diverse, ranging from defense against pathogens in humans and plants, to mammalian organ development and fungal morphogenesis and pathogenicity [2-8]. Additionally, IPPs play an important function in cell adaptation to adverse environmental conditions, including a predominant role in maintaining phosphate homeostasis in mammals, plants and fungi [4, 9-19]. Although, IPPs appear to have rather pleiotropic effects, evidence is accumulating that different phenotypes are connected such as fungal virulence and phosphate homeostasis [20]. Importantly, the cellular levels of IPPs are responsive to different extrinsic signals [21, 22] underscoring their role in regulating intracellular processes in response to environmental changes [23, 24].

IPPs regulate cellular processes via distinct modes of action including direct binding to proteins and pyrophosphorylation of a protein at a pre-phosphorylated serine residue [16, 18, 20, 25-28]. Two highly conserved, exclusively eukaryotic enzyme families generate the two IPPs, 5-PP-IP_5_ (IP_7_) and 1,5-(PP)_2_-IP_4_ (IP_8_) through the IP6K and Vip1/PPIP5K kinases, respectively [29-34].

As other members of the Vip1/PPIP5K family, the fission yeast *Schizosaccharomyces pombe* Asp1 protein is a bifunctional enzyme consisting of an N-terminal kinase domain with 1-kinase activity generating the less abundant IPP IP_8_ and a [2Fe-2S]-binding C-terminal pyrophosphatase domain with specific inositol pyrophosphate 1-phosphatase activity (Figure 1A [7, 10, 13, 35-37]. Asp1 was originally discovered as a modulator of the cortical actin cytoskeleton, but since then numerous biological functions have been identified to be regulated by this protein [6, 7, 13, 19, 38-41]. In particular, the dynamics of the microtubule (MT) cytoskeleton in interphase cells, spindle assembly, and the dynamics and correct association of spindle MTs with the duplicated sister chromosomes are regulated by IP_8_ levels in a dose-dependent manner [39]. Cells unable to generate IP_8_ delay entry into anaphase A due to an activated spindle assembly checkpoint (SAC) indicating a defect in the correct binding of MTs to kinetochores, the attachment point on chromosomes, reviewed in [42]. Genetic deactivation of the SAC in IP_8_-deficient cells resulted in significant chromosome missegregation and the generation of aneuploid cells. Intriguingly, higher than wild-type IP_8_ levels had the opposite effect: chromosome biorientation, i.e., correct association of sister chromosomes with MTs was faster than in the wild-type and increased chromosome transmission fidelity of a specific chromosome [39]. Thus, Asp1 kinase activity is required for genome stability and the avoidance of aneuploidy events, i.e., the loss or gain of whole chromosomes, which is a hallmark of cancer cells and found in several neurodegenerative diseases, reviewed in [43, 44]. To better define Asp1-mediated transition through mitosis, we now determined if IP_8_ was required for entry into M-phase and which components of the MT-kinetochore interface other than MTs were subject to regulation by IP_8._ In particular, we analyzed if yeast strains expressing mutant components of the kinetochore were affected by varying IP_8_ levels. The huge macromolecular kinetochore complexes, which are several-fold larger than ribosomes are assembled on centromeric chromatin, which is defined in many organisms via the centromere-specific histone H3 variant, CENP-A [45], reviewed in [46, 47]. Kinetochore composition is conserved and subcomplexes can be roughly classified as inner (close to the centromeric chromatin) and outer (close to/associating with spindle MTs) kinetochore subcomplexes. The 10-component outer subcomplex KMN (NMS in *S. pombe*) is the main platform for attachment of spindle MTs and the number of copies at the kinetochore and kinetochore localization *per se* are cell cycle controlled in higher eucaryotes [48]. Interestingly, it has been shown recently, that this type of dynamic kinetochore composition is also present in *S. pombe* [49]. The role/importance/presence of specific components of the inner subcomplex constitutive centromere-associated network CCAN (in vertebrates/Ctf19 in *S. cerevisiae*/Mis6-Mal2-Sim4 in *S. pombe*) is somewhat variable depending on the organism analyzed. For example, the *S. pombe* Mal2, the *S. cerevisiae* Mcm21, which is part of the COMA subcomplex and human CENP-O which belongs to CENP-OPQRU subcomplex are all members of one protein family. However, Mal2 is an essential protein, while Mcm21 is not and CENP-O requirement depends on the cell type [50-52]. Overall CCAN recruits and is required to maintain the specific histone H3 variant CENP-A, links centromeric chromatin and outer kinetochore, and recent cyro-EM structures of CCAN and Ctf19 demonstrated the importance of this subcomplex in dealing with spindle generated forces reviewed in [47, 53-57]. Importantly, although the name CCAN implies that this complex is present throughout the cell cycle, abundance of CCAN components can vary during the cell cycle [58].

**Figure 1.**
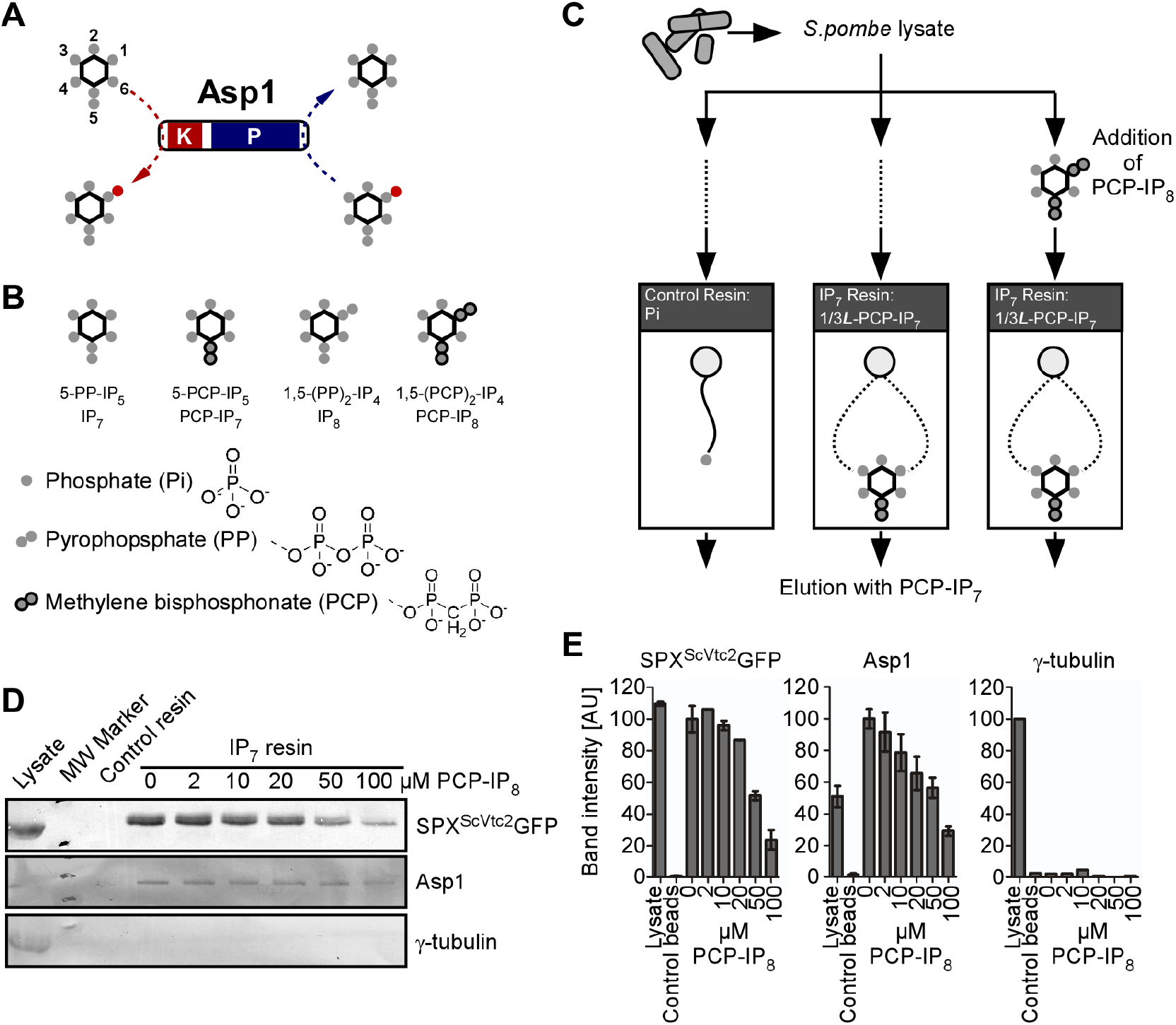
Endogenous Asp1 targets IPPs. **(A)** Schematic representation of the catalytic activities of *S. pombe* Asp1; K, kinase domain; P, pyrophosphatase domain. The *myo*-inositol ring is represented by a hexagon and the position of the phosphates (filled gray circles) on the inositol ring are numbered. The 1-β-phosphate (position C1), which is added and removed by Asp1, is colored red. **(B)** Graphical depiction of IP_7_ and IP_8_ and their corresponding metabolically-stable analogues. **(C)** Experimental overview of the affinity enrichment strategy and reagents utilized for the analysis of *S. pombe* cell lysates. The lysate was aliquoted and a subset of these aliquots was spiked with different concentrations of PCP-IP_8_ prior to incubation with control or IP_7_ resin. The linker utilized for the immobilization of Pi is depicted as a solid line. PCP-IP_7_ is immobilized on beads through a linker attached to the phosphate at either position C1 or C3 of the inositol ring (1/3***L***-PCP-IP_7_). All samples were eluted with PCP-IP_7_. **(D)** Western blot analysis of lysate and eluates from control and IP_7_ resin. The final concentration of PCP-IP_8_ used for each sample is noted. Endogenous proteins were detected with antibodies against Asp1 or γ-tubulin. A GFP antibody was used to detect SPX^ScVtc2^GFP. 1 technical replicate shown. **(E)** Band intensity quantification of proteins analyzed by western blots in (D). Asp1 = 3; SPX^ScVtc2^GFP = 2; γ-tubulin = 1 technical replicate. Arbitrary units (AU).

Our present analysis of kinetochore targeting of fission yeast kinetochore proteins, identifies a new regulator of kinetochore architecture: intracellular IP_8_, which modulates kinetochore presence of specific *S. pombe* CCAN components in a dose-dependent manner.

## Materials and Methods

### Fission Yeast strains and plasmids

All strains are listed in Table S1. Plasmids used are listed in the Table S2. New strains were obtained by crossing the initial strains followed by random spore analysis or tetrad dissection. *S. pombe* strains were grown in rich (YE5S) or minimal medium (MM) with supplements [59]. To repress/de-repress the *nmt1/nmt41* promoter, transformed cells were grown in MM with or without 5 µg/ml thiamine, respectively. This leads to low or high expression of the ORF of interest. For serial dilution patch tests, 10^4^ to 10^1^ cells of the indicated strains transformed with the relevant plasmid were grown under plasmid-selective conditions at the indicated temperatures.

For overexpression of *kcs1*^*+*^, the genomic DNA region encoding *S. pombe kcs1*^*+*^ was amplified from the wildtype yeast strain KG425 and ligated into pJR1-41XL [60] cut with Bsp68I using a blunt ligation approach giving rise to plasmid pUF1489. For the plasmid expressing SPX^ScVtc2^GFP (pUF1577), the genomic DNA fragment encoding the *Saccharomyces cerevisiae* Vtc2 protein, amino acids 1 to 146 (SPX domain) were amplified from a CEN.PK strain and ligated into the XhoI and NotI sites upstream of a nuclear localization signal fused to GFP previously cloned into plasmid pREP3XL [60] (pUF1027).

### Affinity-enrichment of IPP binding proteins

Harvesting and cryogenic lysis of *S. pombe* cells was performed adapting a previously described protocols for *S. cerevisiae* [61]. In short, wild type fission yeast transformed with a plasmid encoding SPX^ScVtc2^GFP under the control of the *nmt1* promoter was grown in 2 l of MM with supplements without thiamine under plasmid selective conditions for 16 h at 25°C. Cultures with an OD_600_ of 0.47 were harvested by centrifugation at 3500 x g for 10 min at 4°C. Cells were collected and washed three times with ice-cold 50mM Tris pH 7.5 with subsequent 5 min centrifugation at 2600 x g at 4°C. Buffer was removed and the pellet was centrifuged two more times removing as much as the buffer as possible after every spin. The yeast pellet was transferred to a syringe with a small spatula and slowly pressed with a plunger to generate short yeast noodles directly into a 50ml falcon tube filled with liquid nitrogen. Liquid nitrogen was decanted and noodles stored at -80°C. Yeast noodles were transferred to a 35ml Steel Mixer Mill Grinding Jar (Retsch) that had been previously cooled down with liquid nitrogen. A cooled 10mm steel ball was placed on top of the noodles and the jar was closed. Jars were placed in a CryoMill (Retsch) and shaken for 3 min at a frequency of 30/s. The jar was cooled down with liquid nitrogen and then shaken again for a total of 5 cycles. The resulting frozen powder was stored at -80°C. Lysis was checked microscopically with thawed powder.

Frozen yeast lysate powder was resuspended in ice-cold lysis buffer (50M Tris-HCl, pH 7.4, 150mM NaCl, 0.05% Triton X-100, 1x Roche cOmplete Proteinase inhibitor cocktail) to a concentration of 6µg/µl and rotated for 10 min at 4°C before centrifugation at 3000 x g for 10min at 4°C. The lysate was aliquoted and specified samples were spiked with different concentrations of PCP-IP_8_ synthesized as in [62]. Lysates were then incubated with 100 µl equilibrated PCP-IP_7_ beads generated as in [63] with constant rotation at 4°C. Samples were then centrifuged at 2000 x g for 2 min and the supernatant was removed. Beads were washed three times with 1 ml lysis buffer for 5 min with rotation and centrifugation at 2000 x g for 2 min. Beads were then incubated with 100µl of lysis buffer containing 5mM PCP-IP_7_ for 30 min under constant rotation at 4°C. Finally, samples were centrifuged and supernatant was collected as the elution fraction, which was further analyzed by western blotting using anti-GFP, anti-Asp1 and anti-γ-tubulin antibodies (Table S2). Quantification of band intensity was performed with Image Lab (Bio-Rad).

### Microscopy

Immunofluorescence microscopy of fixed *S. pombe* cells was carried out as described [64]. Transformed cells were grown in plasmid-selective MM without thiamine overnight prior to analysis. First antibody used: monoclonal α-tubulin antibody TAT1 [65]. Secondary antibody: α-mouse Alexa Fluor®488 (1:200; Thermo Fisher Scientific). DNA was stained with 4,6-diamidino-2-phenylindole (DAPI). Phenotypes were determined and counted visually using a Zeiss Axiovert200 fluorescence microscope. Pictures shown in Figure 3D were taken using a Nikon Eklipse Ti microscope.

For DAPI staining of yeast cells, liquid cultures were grown at 25°C overnight in YE5S. Cultures were then divided and grown for 6 hours at either 25°C or 33°C. 1 ml of each culture was centrifuged at 3000 x g, and fixed in 70% ice-cold ethanol. Cells were then washed 1x with PBS and stained with 100 ng/ml DAPI. Image acquisition for Figure 2H was done with a Zeiss Axiovert 200 fluorescence microscope (Carl Zeiss) using a 63x objective with a charge-coupled-device (CCD) camera (IEEE1394-Based Digital Camera Orca-ER 1394; Hamamatsu, Herrsching, Germany) and image editing and analysis was done with ImageJ 1.47v (National Institutes of Health).

**Figure 2.**
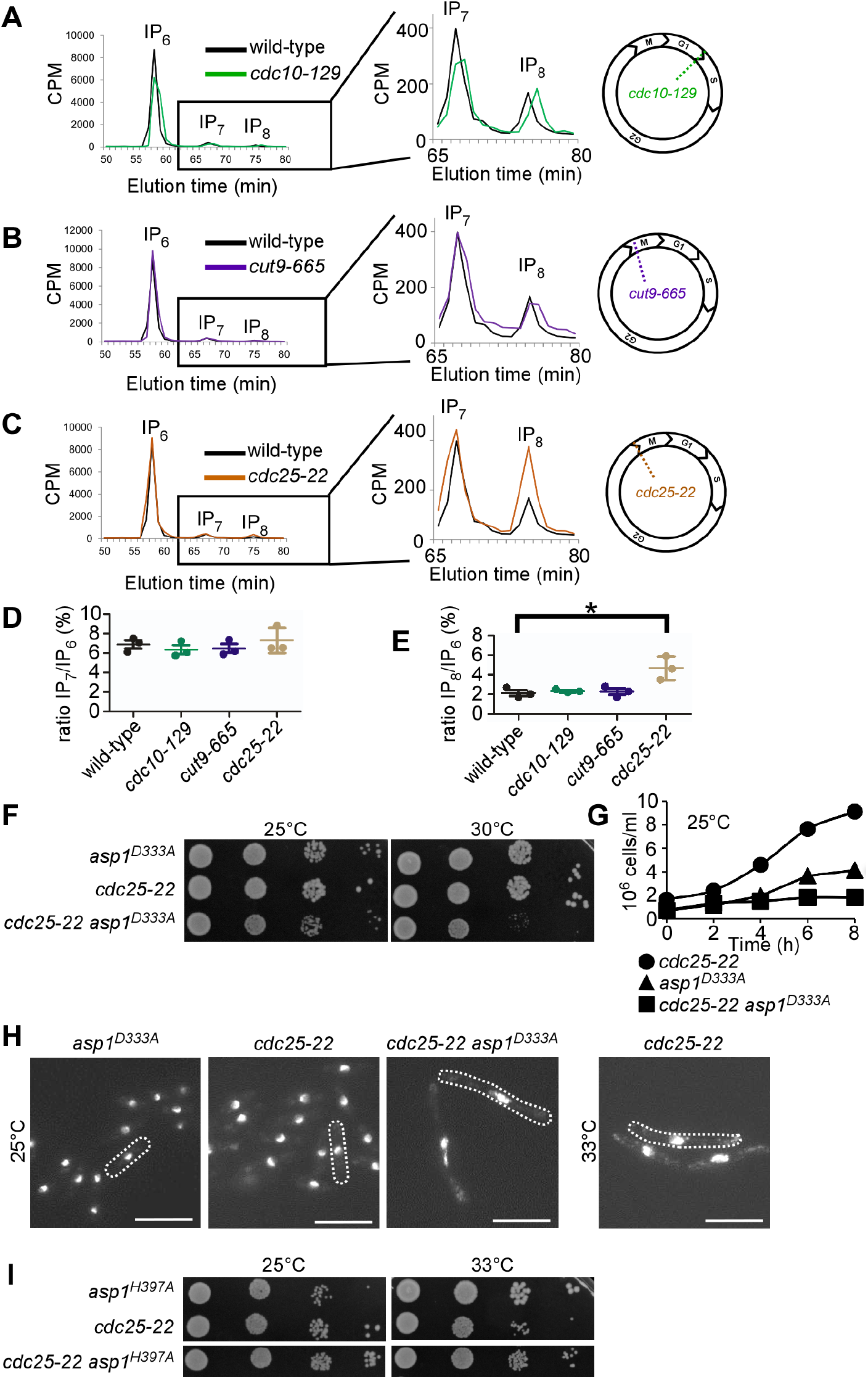
IP_8_ is cell-cycle regulated and required for mitotic entry. **(A-C)** Typical HPLC profiles of soluble inositol phosphates and pyrophosphates extracted from cells arrested in **(A)** G1 (*cdc10-129)*; **(B)** metaphase-anaphase A transition (*cut9-665)* and **(C)** G2/M boundary (*cdc25-22)* compared to wild-type controls treated as the *cdc* strains. Boxed regions define the magnification of the IP_7_ and IP_8_ peaks shown on the right side. A schematic representation indicating where in the cell-cycle the indicated mutant arrests is shown in the right most panels. Wild-type and temperature-sensitive *cdc* strains were labeled with [^3^H]inositol and pre-grown at 25°C before a shift to 36°C for 6 hours prior to the soluble inositol extraction. Counts per minute (CPM). **(D)** Quantification of the IP_7_ levels relative to IP_6_. Wildtype = 6.88 ± 0.72; *cdc10-129* = 6.35 ± 0.79; *cut9-665* = 6.48 ± 0.78; *cut25-22* = 7.29 ± 1.31 **(E)** Quantification of the IP_8_ levels relative to IP_6_. Wildtype = 2.12 ± 0.5; *cdc10-129* = 2.34 ± 0.14; *cut9-665* = 2.26 ± 0.57; *cdc25-22*: 4.68 ± 1.18. For (D) and (E), mean and SD of n = 3 are shown. *, p = 0.0258 using a t-test. **(F)** Serial dilution patch test (10^4^-10^1^ cells) of indicated strains grown for 4 (30 °C) or 6 days (25 °C) on YE5S plates. **(G)** Growth rates of indicated strains grown in liquid YE5S at 25°C. **(H)** Photomicrographs of fixed cells stained with DAPI to reveal the nucleus. Dotted lines indicate the cell periphery. Scale bars, 10 µm. **(I)** Serial dilution patch test (10^4^-10^1^ cells) of indicated strains grown at the indicated temperatures on YE5S plates.

For live-cell imaging, cells were grown overnight at the indicated temperature in sterile-filtered MM with supplements (LFM: live fluorescence media). Microscopy slides were prepared by patching cells on agarose pads made of LFM containing 2% agarose and were sealed with VALAP (vaseline, paraffin and lanolin in a 1:1:1 ratio) [66]. A Zeiss spinning-disk confocal microscope equipped with a Rolera EM-C^2^ (QImaging) camera (or an Axiocam 702 mono camera or a Zeiss LSM 880 Airyscan microscope with GaAsP, PMT and T-PMT detectors were used for imaging (CAi; HHU Düsseldorf). LSM: Figure 4B. Spinning disk: Figures 5C, 6B, 6C, S3A and S3B. Imaging and analysis were performed with Zen2012 and AxioVision software. Image processing was done with ImageJ and Canvas 14. In all figures clippings of Maximum Intensity Projection (MIP) images are shown. The contrast and brightness settings were chosen equally within data sets.

**Figure 3.**
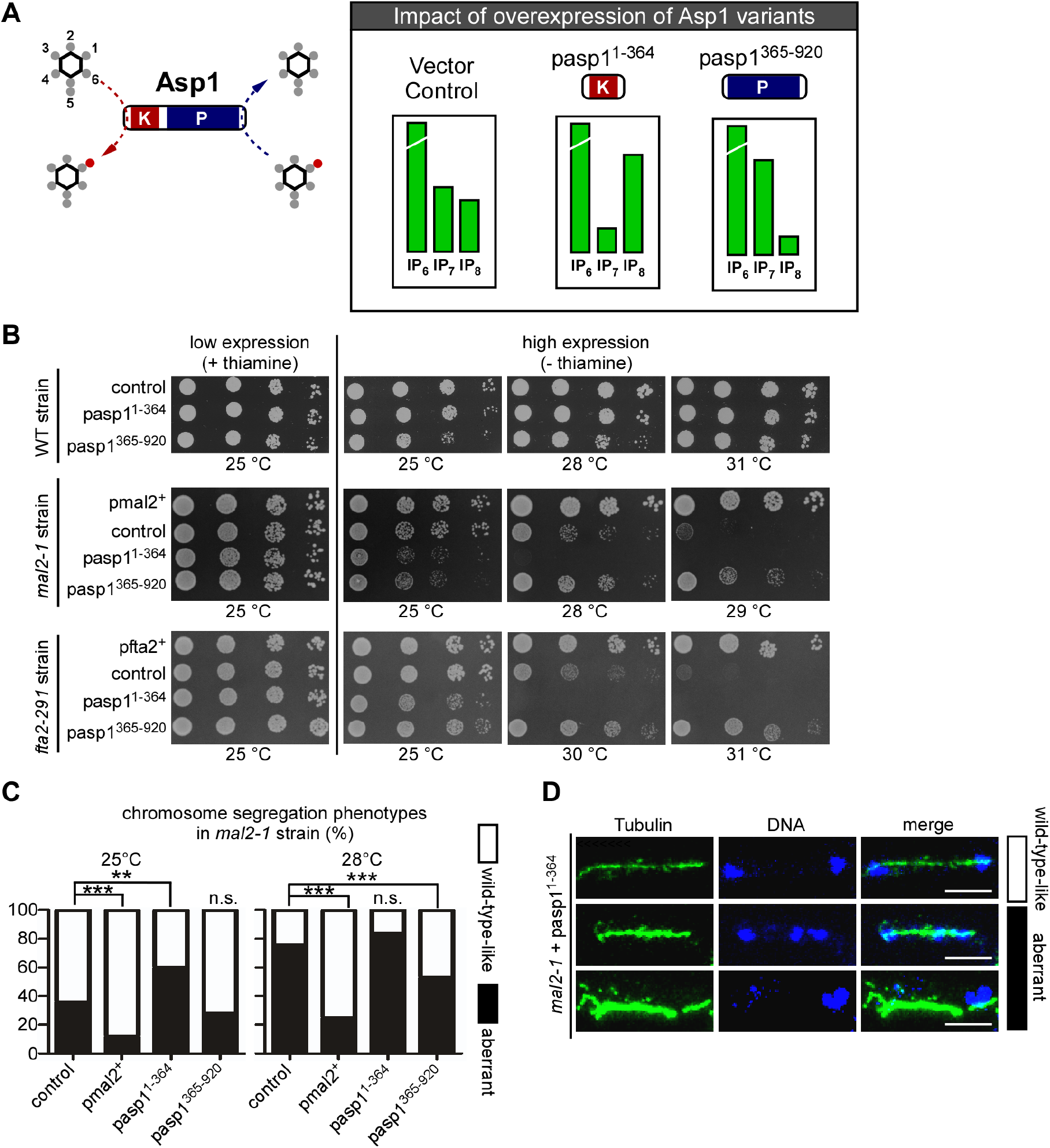
Alteration of IP_8_ levels affect the temperature-sensitive growth phenotype of CCAN kinetochore mutant strains. **(A)** Schematic representation of the impact in IP_7_ and IP_8_ levels upon expression of the Asp1 kinase (Asp1^1-364^) and pyrophosphatase (Asp1^365-920^) variants [36, 37]. **(B)** Serial dilution patch test (10^4^-10^1^ cells) of wild-type (WT), *mal2-1* and *fta2-29*1 strains transformed with the indicated plasmids and grown at the indicated temperature for 4-8 days depending on the incubation temperature. pmal2^+^; plasmid with wild-type *mal2*^*+*^ ORF expressed via the *nmt1* promoter. *nmt1* transcribed genes show low expression in the presence of thiamine and high expression when no thiamine is present. pasp1^1-364^ and pasp1^365-920^, plasmids with *nmt1* driven expression of the Asp1 kinase or pyrophosphatase domains, respectively. One of n = 3 shown. **(C)** Quantification of chromosome segregation phenotypes observed in a *mal2-1* strain transformed with the indicated plasmids. 25 °C: **, p = 0.0011 ***; p < 0.0001. 28 °C: ***, p < 0.0001 (*mal2*^*+*^); ***, p = 0.0007 (asp1^365-920^), n.s., not significant. Fisher’s exact test. n = 100 cells per plasmid. One of n = 2 shown. **(D)** Immunofluorescence of fixed *mal2-1* cells transformed with pasp1^1-364^ and grown at 25 °C. Green, α-TAT1 (tubulin); blue, DAPI staining. Scale bars, 5 µm.

**Figure 4.**
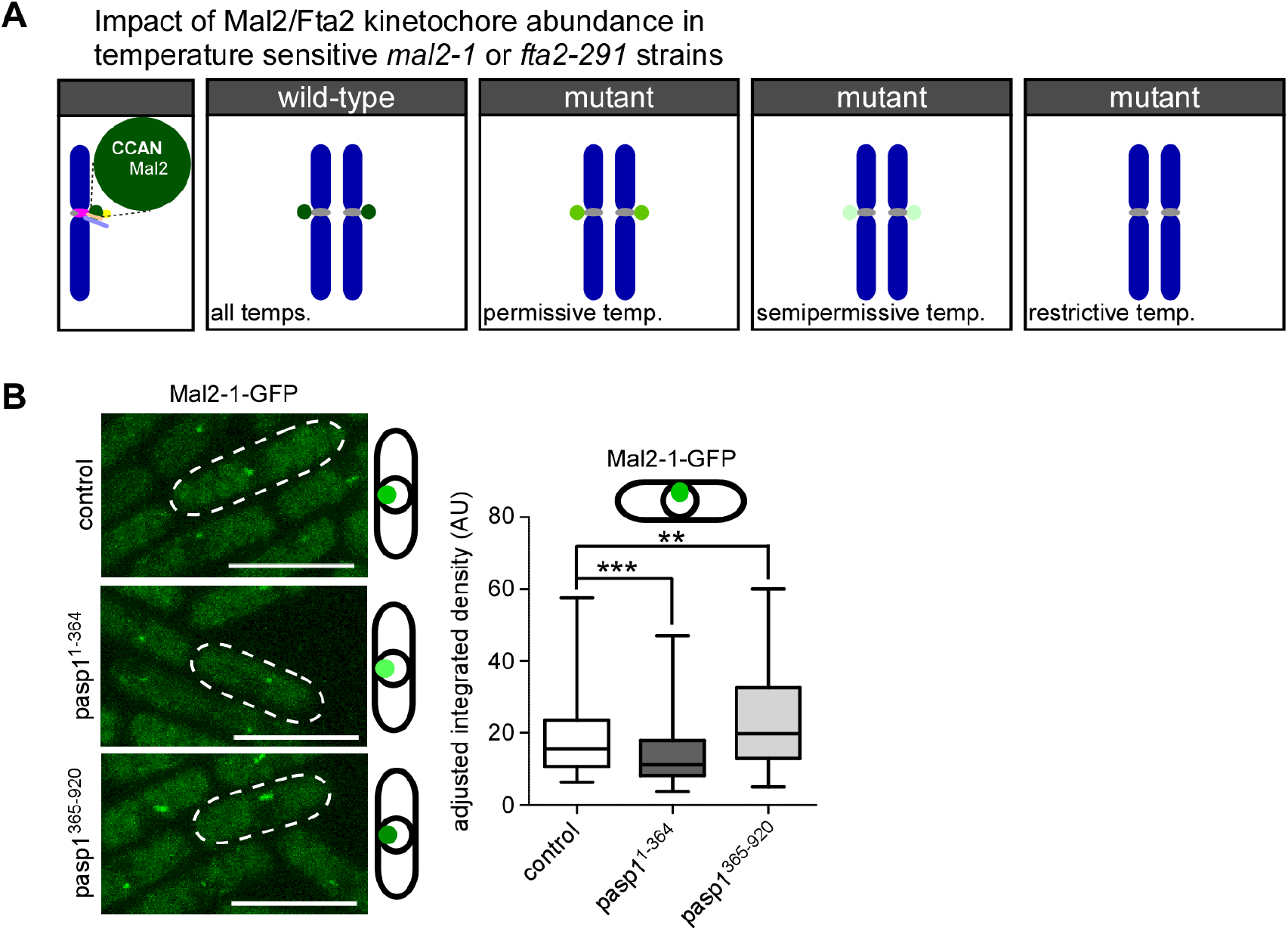
Negative correlation between IP_8_ levels and Mal2-1-GFP kinetochore fluorescence. **(A)** Diagrammatic representation to show that Mal2-1-GFP kinetochore targeting is temperature dependent [91]. The lighter the shade of green the less Mal2-1-GFP is at the kinetochore. **(B)** Left: Live cell images of the indicated *mal2-1-gfp* transformants grown at 25 °C. Scale bar, 10 µm. Right: Quantification of Mal2-1-GFP fluorescence signals: Mean and SD: vector = 18.42 AU ± 9.7; *asp1*^*1-364*^ = 14.53 AU ± 6.39; *asp1*^*365-920*^ = 22.77 AU ± 12.17. Number of kinetochore signals counted: control: = 162; *asp1*^*1-364*^ = 160; *asp1*^*365-920*^ = 140; ***, *p* < 0.0001; **, *p* = 0.0019; Mann-Whitney U-Test. One of n = 2 or 3 sets shown.

**Figure 5.**
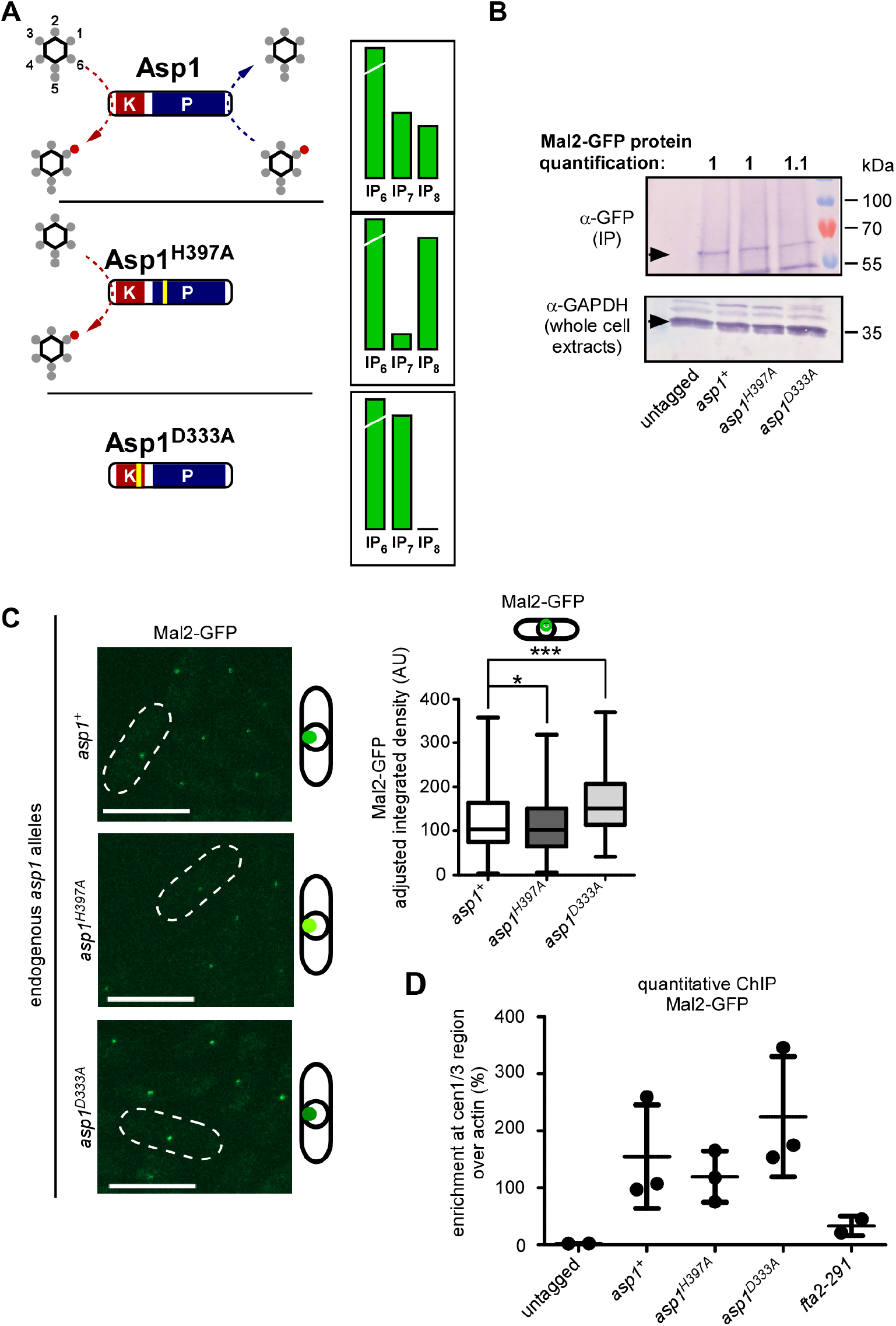
Kinetochore-targeting of wild-type Mal2 is subject to IP_8_ levels. **(A)** Schematic representation of the three endogenous Asp1 variants and their impact on IP_7_ and IP_8_ levels [36]. Note that in the case of Asp1^D333A^, no IP_8_ is generated and thus, no IP_8_ is being hydrolyzed, although the pyrophosphatase domain is intact. **(B)** Western blot analysis of Mal2-GFP protein levels in specified *asp1*-variant strains utilizing indicated antibodies. To detect Mal2-GFP, an immunoprecipitation with anti-GFP antibodies was performed prior to western blot analysis. Quantification of fold change of Mal2-GFP bands normalized to GAPDH are shown above the blot. One of n = 2 sets shown. **(C)** Left: Live-cell images of the indicated *asp1*-variant strains endogenously expressing *mal2*^*+*^*-gfp* grown at 25°C. Scale bars, 10 µm. Right: Quantification of Mal2-GFP fluorescence signals in each strain. Mean and SD: *asp1*^*+*^= 122.5 AU ± 63.67; *asp1*^*H397A*^ = 110.8 AU ± 58.43; *asp1*^*D333A*^= 163.8 AU ± 65.53. Number of kinetochore signals counted, *asp1*^*+*^ n = 381; *asp1*^*H397A*^ n = 381; *asp1*^*D333A*^ n = 380. *, *p* = 0.0195; ***, *p* < 0.0001; Mann-Whitney U-Test. Average of n = 3 shown. **(D)** qChIP analysis for the strains imaged in (C). Shown is enrichment of cen1/3 DNA relative to the *act1*^*+*^ locus. Mean and SD shown: untagged (wildtype strain) = 1.99 ± 0.32; *asp1*^*+*^ = 154.67 ± 90.99; *asp1*^*H397A*^ = 119.55 ± 45.01; *asp1*^*D333A*^ = 224.83 ± 105.56; *fta2-291* = 33.03 ± 17.06. Untagged and *fta2-291* strains, n = 2. *asp1*^*+*^, *asp1*^*D333A*^, *asp1*^*H397A*^ strains n = 3 biological replicates with 2-4 technical replicates each.

### Protein extraction, IP and Western Blot analysis of Mal2-GFP

For Western Blot analysis of Mal2 strains were grown overnight in YE5S. 5×10^8^ -1×10^9^ cells were washed once with 5 ml STOP buffer (0.9% NaCl, 1 mM NaN_3_, 10 mM EDTA, and 50 mM NaF). Cells were resuspended in 500 μl HB15 lysis-buffer (25 mM MOPS, 60 mM ß-glycerophosphate, 15 mM p-nitrophenyl phosphate, 15 mM MgCl_2_, 15 mM EGTA, 1 mM DTT, 0.1 mM sodium orthovanadate, 1% TritonX100, 1 mM PMSF and cOmplete protease inhibitor (Roche Diagnostics)) and lysed using glass beads [67]. The protein extract was cleared twice by centrifugation at 13000 rpm for 30 min at 4 °C. For immunoprecipitation, 250 μl of each sample were incubated on ice for 1 h with 50 μl of α-GFP µMACS beads (Miltenyi Biotec). Immunoprecipitates were isolated using the μMACS GFP isolation kit. Columns were equilibrated with 200 µl HB15 lysis-buffer before use. Columns were washed 8x with 200 µl HB15 lysis-buffer and proteins isolated in 2x 50 µl elution buffer. Eluates or whole cell extracts were resolved on 10% SDS-gels before blotting. If the expected proteins were of different molecular weights membranes were cut to detect different proteins on one membrane. Antibodies used in Western blot analysis: monoclonal α-GFP (1:1000; mouse; Roche), α-GAPDH (1:3000; mouse; Sigma Aldrich) and α-γ-Tubulin (1:10.000; mouse; Sigma Aldrich).

To quantify protein levels, ImageJ 1.44 (NIH) was used to measure the intensity of the protein bands in question. Protein amounts were normalized to the control signal for GAPDH or γ-Tubulin. The value was set to 1 for the wildtype (Figure 5B).

### qChIP

Chromatin immunoprecipitation (ChIP) with *mal2*^*+*^*-gfp S. pombe* strains was performed as follows [68-70]: 200 ml overnight cultures (sterile-filtered MM with supplements) with an OD_600_ of 0.4-0.8 grown at 25 °C from liquid pre-cultures were used. Cells were fixed with 3% paraformaldehyde for 30 min at 25 °C followed by washing twice with 20 ml cold 1x PBS and spheroplasted in 20 ml PEMS (100 mM Pipes, pH 7,1 mM EDTA, 1 mM MgCl_2_, 1.2 M sorbitol) with 50 mg/ml Lallzyme at 37 °C for 30-45 min. Samples were washed twice in 10 ml PEMS and resuspended in 1 ml PEMS. The sample was divided onto two fresh tubes (no antibody control and IP sample). Cells were pelleted and pellets frozen at -20 °C until further use. Pellets were resuspended in 400 µl cold lysis buffer (50 mM HEPES-KOH pH7.5, 140 mM NaCl, 1 mM EDTA, 1 % Triton-X-100, 0.1% sodium deoxycholate) supplemented with 1:100 cOmplete protease inhibitor cocktail and 2 mM PMSF (added shortly before use). Samples were sonicated twice for 6 sec at 10 %. Supernatant was cleared via two 15 000 rpm centrifugation at 4°C for 5 and 10 min. 25 µl of protein A agarose were added to each sample and tubes were incubated on a wheel at 4 °C for 1-2 h. Supernatant was cleared by centrifugation at 8000 rpm for 5 min at 4 °C. 40 µl of each sample were frozen for later processing as input control. 2 µl α-GFP antibody and 25 µl protein A agarose was added to the remaining lysates and precipitation was performed at 4 °C on a wheel overnight. The protein A agarose was pelleted via centrifugation at 8000 rpm for 5 min at 4 °C. Protein A agarose was washed with 1 ml of the following buffers followed by 5 min spinning on a wheel and 2 min centrifugation at 8000 rpm at each step: 1) lysis buffer, 2) lysis buffer with 0.5 M NaCl, 3) wash buffer (10 mM Tris/HCl pH: 8, 250 mM LiCl, 1 mM EDTA, 0.5 % NP-40, 0.5 % sodium deoxycholate) and 4) TE-buffer. 250 µl TES (TE-buffer + 1 % SDS) were added to the agarose and 210 µl to the formerly frozen 40 µl input control and samples incubated at 65 °C overnight in a water bath. Supernatant was cleared by 1 min centrifugation at 8000 rpm. 450 µl TE supplemented with 30 µl freshly prepared Proteinase K (10 mg/ml) were added to each tube followed by incubation for 4 h at 37 °C while shaking. Afterwards phenol:chloroform and chloroform extraction were performed. DNA was precipitated by addition of 1:10 v/v 3 M NaAc pH 5.5 and 2.5:1 v/v 96 % EtOH and incubation on dry ice for 1 h. Samples were centrifuged for 30 min at 15.000 rpm at 4 °C and pellets air-dried. IP samples were resuspended in 30 µl TE-buffer and inputs in 300 µl TE-buffer. Samples were stored at -20 °C and 5 µl of 1:10-1:50 dilutions (equal within one experimental set and decided after analysis of the DNA amount in the input sample on an agarose gel) used for qPCR. For qPCR reactions the GoTaq qPCR Master Mix (Promega) was used following manufacturers instructions. For each sample qPCR was performed with two primer sets. The first set amplifies a region within the central region of centromeres 1 and 3 (cen1/3). The second set amplifies a fragment within the *act1*^*+*^ locus on chromosome 2 (actin) [71, 72]. The oligonucleotides are listed in Table S2.

### [^3^H]inositol labeling and HPLC analysis

For [^3^H]inositol labeling and soluble inositol extraction *S. pombe* cultures were grown overnight in MM containing all supplements and 10 μM inositol at 25 °C [36]. For cell-cycle arrest experiments cultures were then diluted to an OD_600_ of 0.05 in 5 ml MM supplemented with 10 μM inositol and 5 μCi/ml of [^3^H]inositol (10-25Ci/mmol) and incubated at 25 °C. After 15 h each culture was shifted to 36 °C for 6 h. For non-shifted cultures the cells were kept at 25 °C.

Inositol polyphosphates were extracted from the labeled cultures. Cells were centrifuged for 2 min at 2000 rpm at room temperature, washed once with 1 ml H_2_O and transferred to 1.5 ml tubes. Cells were resuspended in 200 µl 1 M Perchloric acid + 5 mM EDTA (freshly added). ∼1/2 PCR tube of glass beads was added and the cells lysed in a gene disrupter for 5 min at 4 °C. The supernatant was cleared by 5 min centrifugation at 14.000 rpm at 4 °C. The cell pellet was kept at this step; later 1 ml 0.1 % NaOH + 0.1 % SDS are added to measure total amount of radioactivity later; samples were rotated on a wheel at room temperature overnight. 45 µl of 1 M K_2_CO_3_ + 5 mM EDTA (freshly added) were added to the supernatants and the tubes kept on ice for 2 h with their lids open (tubes were flicked gently every 15-30 min; CO_2_ bubbles form and evaporate). The pH of the samples was determined and had to be between 6 and 8. Samples were centrifuged for 5 min at 14.000 rpm at 4 °C and supernatants transferred to a new tube and kept at 4 °C until HPLC analysis. Inositol polyphosphates were then resolved by strong anion-exchange SAX-HPLC (using a PartiSphere SAX 4.6×125mm column; Hichrom). 1 ml fractions were collected each minute over 80 min. 4 ml of Ultima-Flo AP scintillation cocktail were added to each fraction followed by vigorous mixing. All fractions were analyzed via scintillation counting [73].

### Quantification of IP_8_/IP_6_ ratios

The CPM values measured were used to generate the graphs shown in Figure 2. To calculate IPP ratios we first determined the background signal by measuring CPM in the last two fractions of the entire HPLC run. This amount was subtracted from all the values used for quantification of peaks. The values that were part of the peaks for IP_6_, IP_7_ and IP_8_ were summed up for calculation. The ratio is given as percentage of the indicated IPP when compared to IP_6_.

## Results

### IP_8_ is targeted by endogenous Asp1 and its level is cell-cycle controlled

We and others have described the catalytic properties of recombinant Asp1 purified from bacterial expression systems [7, 10, 31, 36, 74]. In these *in vitro* assays, Asp1 kinase and pyrophosphatase domains can independently interact with IPP substrates and perform their catalytic functions at position C1 of the inositol ring before releasing their products (shown diagrammatically in Figure 1A). To determine the interaction of endogenous wild-type Asp1 protein with its IP_7_ substrate, we performed an affinity-based enrichment utilizing the metabolically stable IP_7_ analog 5-PCP-IP_5_ (PCP-IP_7_) (Figure 1B) immobilized to agarose beads (IP_7_ resin). As control beads, the same beads and linker attached to a negatively charged phosphate group were used (control resin) (Figure 1C). As a positive control for IP_7_ binding, we used the SPX domain of the *S. cerevisiae* Vtc2 protein, a well-characterized IPP-binding module [16] fused to GFP (SPX^ScVtc2^-GFP). Additionally, in order to observe if IP_8_ was able to compete for binding to IP_7_, part of the *S. pombe* lysate was first incubated with different amounts of the IP_8_ analogue 1,5-(PCP)_2_-IP_4_ (PCP-IP_8_) before exposing it to the IP_7_ resin.

Clarified lysates of a wild-type fission yeast strain (i.e., expressing endogenous *asp1*^*+*^), transformed with a plasmid encoding SPX^ScVtc2^-GFP were spiked with 0, 2, 10, 20, 50 or 100µM of PCP-IP_8_ and applied to control or IP_7_ resin, followed by washes with binding buffer and elution with 10mM PCP-IP_7_. Eluates were analyzed by western blotting using GFP, Asp1 and γ-tubulin antibodies (Figure 1D-E). Both SPX^ScVtc2^-GFP and Asp1 bound to the IP_7_ resin with great specificity when compared to control beads, similar to what has been found for the human PPIP5K family members [75]. However, pre-incubation of lysates with PCP-IP_8_ hindered binding of SPX^ScVtc2^-GFP to the IP_7_ resin, confirming that SPX domains can bind IP_8_ [76]. Interestingly, binding of the Asp1 protein to IP_7_ resin was blocked by PCP-IP_8_ presence in a dose-dependent manner, demonstrating that endogenous Asp1 bifunctional protein has IPP binding pockets able to bind IP_7_ and IP_8_.

To decipher the numerous biological roles of *S. pombe* Asp1, we previously used strains expressing Asp1 mutant variants which gave rise to altered intracellular IPP levels [6, 7, 36, 39]. This was accomplished by mutations in either the kinase - or pyrophosphatase domain of the Asp1 protein [36] and this decrease/increase of the IP_8_ output had consequences for cellular functions regulated by IP_8_ in fission yeast in a dose dependent manner [6, 7, 36, 39]. However, it was unclear if physiological modulations of this bifunctional enzyme activity occur especially as Asp1 protein levels appear to remain constant during the cell cycle [77]. Thus, we assayed if the IP_8_ levels changed during the *S. pombe* cell-cycle. The inositol polyphosphate species were determined in three temperature-sensitive cell cycle mutant strains, *cdc10-129, cdc25-22* and *cut9-665*, which arrest at the restrictive temperature in G1, at the G2/M transition and before anaphase A, respectively [78-81]. Mutant cells were pre-grown at the permissive temperature of 25 °C followed by 6 hours incubation at the restrictive temperature of 36 °C. The cell cycle arrest was monitored by microscopy as arrested *cdc10-129* and *cdc25-22* cells are elongated and arrested *cut9-665* cells show highly condensed chromosomes. To quantify the abundance of IP_7_ and IP_8_ in these strains, its precursor IP_6_ was used as a reference. The ratio of IP_7_/IP_6_ was not influenced by the cell-cycle stage in any of the three mutant strains tested (Figure 2A-C; quantification in 2D). However, while the IP_8_/IP_6_-ratios in cells arrested in G1 (*cdc10-129*) and at the metaphase/anaphase A transition (*cut9-665*) were comparable to non-arrested wild-type cells incubated at 36 °C (Figure 2E), the IP_8_/IP_6-_ratio was significantly increased in *cdc25-22* cells (Figure 2C and quantification in 2E). Mitosis is universally initiated by the activation of Cdk1 protein kinases, which in turn are controlled by Cdc25 phosphatases [82]. Thus, *cdc25-22* mutant cells arrest in late G2 at the transition to M-phase and at this cell cycle stage the IP_8_/IP_6-_ratio was increased. Such an increase was not observed at the permissive temperature of 25 °C, when these cells can progress through the cell cycle (Figure S1 A-C). As the increase at the G2/M transition might indicate a particular relevance of IP_8_ at this cell cycle stage, we tested if IP_8_ was required for mitotic entry. We therefore generated a *cdc25-22 asp1*^*D333A*^ double mutant strain. The Asp1^D333A^ mutant protein carries a single amino acid change in the catalytic domain of the kinase which leads to a kinase-dead version and no cellular IP_8_ [36] (Figure 5A). Patch test analysis of the double mutant and parental strains showed that the *cdc25-22 asp1*^*D333A*^ strain grew slightly slower on solid media at 25 °C than the single mutant *asp1*^*D333A*^ strain and had a clearly reduced growth at 30°C compared to the parental strains (Figure 2F). In addition, the generation time of the double mutant at 25 °C was nearly 6 hours, twice as long as that of the parental strains, which resulted in a very shallow growth curve in liquid media when compared to single mutant strains (Figure 2G). Microscopic analysis of double mutant cells grown at 25 °C in liquid revealed that these cells were highly elongated with a stretched-out nucleus (Figure 2H). This phenotype is the hallmark of *cdc25-22* cells incubated at semi-permissive/restrictive temperatures (Figure 2H, right most panel) [83]. Intriguingly, when *cdc25-22* cells have more IP_8_, i.e., in an *asp1*^*H397A*^ background (Figure 5A diagrammatically depicts IPP levels in asp1^H397A^ strain) the reduced growth phenotype seen for the *cdc25-22* mutant grown at the semi-permissive temperature of 33 °C is rescued (Figure 2I). We conclude that IP_8_ is required for entry into mitosis.

### Alteration of intracellular IP_8_-levels can rescue non-growth of *S. pombe* CCAN kinetochore mutant strains

Our finding that Asp1 kinase activity was required for entry into mitosis is in accordance with previous work where we have shown that chromosome segregation fidelity is modulated by Asp1 kinase function [39]. Specifically, we established that bipolar spindle formation/function and the accuracy of chromosome transmission required intracellular IP_8_ in a dose-dependent manner. To determine if modulation of spindle dynamics was the sole target of Asp1 regulation during mitosis we now tested if the conductor of mitosis, the kinetochore, was also subject to control by Asp1 made IPPs. We assayed if expression of either the Asp1-kinase (Asp1^1-364^) or the Asp1-pyrophosphatase (Asp1^365-920^) altered growth of specific kinetochore mutant strains. As shown diagrammatically in Figure 3A, plasmid-borne expression of *asp1*^*1-364*^ increases intracellular IP_8_, while *asp1*^*365-920*^ expression reduces (but does not eliminate) IP_8_ levels [36]. *S. pombe* kinetochores complexes consist of the same core modules found in human and *S. cerevisiae* such as KMN(human)/NMS(*S. pombe*) and CCAN(human)/Ctf19(*S. cerevisiae*)/Mis6-Mal2-Sim4(*S. pombe*) [84-88]. Expression of either *asp1*^*1-364*^ or *asp1*^*365-920*^ from a plasmid had no or only a moderate effect on the growth of 4 temperature sensitive *S. pombe* KMN mutant strains tested (Figure S2A). However, when these Asp1 variants were expressed in 5 strains with mutant *S. pombe* CCAN components, we observed a strong effect between growth at higher temperatures and intracellular IP_8_ levels for 4 mutant strains (Figure 3B, Figure S2A-B). For example, the *mal2-1* strain was unable to grow at the semi-permissive temperature of 28 °C when *asp1*^*1-364*^ expression resulted in higher intracellular IP_8_, while its non-growth phenotype at 29 °C was suppressed by the expression of *asp1*^*365-920*^ (lower than wild-type IP_8_ level) (Figure 3B, middle panels). Similarly, the temperature-sensitivity of the *fta2-291* and *mis6-302* mutant strains was increased in cells with higher than wild-type levels of IP_8_ and decreased in cells with reduced IP_8_ level (Figure 3B, bottom panels; Figure S2B). Interestingly, *mis15-68* transformants showed the opposite behavior, as higher than wild-type IP_8_ levels rescued the non-growth phenotype at the non-permissive temperature of 33 °C while decreasing intracellular IP_8_ had the opposite effect (Figure S2B). Thus, the temperature-sensitive growth phenotype of mutant *S. pombe* CCAN components can be altered by varying intracellular IP_8_ levels.

To determine the molecular basis of this phenotype, we used the *mal2-1* mutant strain for further analysis. We assayed chromosome segregation in *mal2-1* transformants grown at 25 or 28 °C by microscopic analysis of fixed mitotic cells. Although *mal2-1* cells are able to grow at 25 °C, the fidelity of chromosome transmission is already decreased at this temperature and 36% of mitotic cells show unequal segregation of chromosomes due to non-disjunction of sister chromosomes as analyzed by staining fixed cells with DAPI and anti-tubulin antibody [50, 69] (Figure 3C, examples of this phenotype shown in Figure 3D). The aberrant phenotype at 25 °C of unequally/partially segregated chromatin on an elongating spindle was reduced to 12 % and 28% in mitotic cell populations expressing plasmid-encoded wild-type *mal2*^*+*^ or *asp1*^*365-920*^, respectively. Expression of *asp1*^*1-364*^ significantly increased the chromosome missegregation phenotype at 25 ° C to 60 % (Figure. 3C). A similar pattern was observed for transformed *mal2-1* cells incubated at 28 °C (Figure 3C), revealing an inverse relationship between chromosome segregation fidelity of *mal2-1* cells and IP_8_-levels.

### Kinetochore-targeting of the mutant Mal2-1 protein is subject to IP_8_ levels

To analyze if kinetochore-targeting of Mal2-1-GFP was dependent on cellular IP_8_-levels, we performed live-cell fluorescence microscopy of endogenous *mal2-1-gfp* interphase cells transformed with plasmids expressing either *asp1*^*1-364*^ or *asp1*^*365-920*^. Growth behavior of this strain transformed with pasp1^1-364^ or pasp1^365-920^ was comparable to the non-tagged *mal2-1* strain (Figure S2C). In interphase cells, the centromeres are clustered at the spindle pole body due to a linker complex and thus fluorescent kinetochore proteins are seen as a single dot-like signal in the nucleus [89, 90]. The Mal2-1-GFP protein is present at the kinetochore at 25 °C, albeit in reduced amounts compared to wild-type Mal2-GFP. With increasing temperatures, the Mal2-1-GFP signal is further reduced or absent (Figure 4A) [50, 69, 91]. Expression of *asp1*^*1-364*^ led to a significant reduction of the Mal2-1-GFP kinetochore signal, while expression of *asp1*^*365-920*^ increased kinetochore-presence of Mal2-1-GFP (Figure 4B). Thus, kinetochore targeting of the mutant Mal2-1 protein is modulated by the intracellular IP_8_ levels.

### Wild-type Mal2 kinetochore-targeting is modulated by intracellular IP_8_-levels

To assess, if IP_8_ kinetochore-targeting also applied to the wild-type Mal2 protein, we determined endogenous Mal2-GFP localization in cells with twice as much IP_8_ as a wild-type strain (Asp1^H397A^) and cells with no IP_8_ (Asp1^D333A^), and compared them to a wild-type control (diagrammatically shown in Figure 5A; based on data from [36]. The total Mal2-GFP protein levels were unaffected by changes in intracellular IP_8_ levels (Figure 5B). However, as had been observed for the mutant Mal2-1 protein, we found that Mal2-GFP kinetochore targeting was also controlled by IP_8_-levels (Figure 5C). Higher than wild-type intracellular IP_8_ significantly reduced Mal2-GFP kinetochore signal, while absence of IP_8_ had the opposite effect (Figure 5C).

Next, we performed a quantitative chromatin immunoprecipitation (qChIP), in which Mal2-GFP was cross-linked to centromeric DNA and the abundance of these DNA regions (cen1/3) over a non-centromere region (actin) was quantified as a measure of kinetochore localization [68-70, 72, 92]. The mutant *fta2-291* strain was used as a negative control for Mal2-GFP kinetochore localization as Mal2 and Fta2 kinetochore localization is interdependent and their orthologs exist as a heterodimer [58, 91, 93-95]. As described previously, the presence of the mutant Fta2-291 protein led to massively reduced Mal2 at the kinetochore (Figure 5D; [91]). Compared to cells with physiological IP_8_-levels (*asp1*^*+*^ strain), the enrichment of cen1/3 over actin increased by ∼45 % in cells without IP_8_ (*asp1*^*D333A*^ strain) and decreased by ∼20 % in cells with more than wild-type IP_8_ (*asp1*^*H397A*^ strain) (Figure 5D). Thus, kinetochore-targeting of the wild-type Mal2 protein is dependent on intracellular IP_8_-levels.

Finally, as the ts phenotype of the *fta2-291* strain was affected in a very similar way as the *mal2-1* strain by altered IP_8_ levels (Figure 3B), we determined if kinetochore targeting of the wild-type Fta2 protein was affected by IP_8_ levels. Fta2-GFP kinetochore-targeting was also IP_8_ regulated as it significantly increased in IP_8_-less cells and decreased in cells with higher-than-wild-type IP_8_ (Figure S3A).

The above microscopic analysis of kinetochore localization of Mal2-1-GFP, Mal2-GFP and Fta2-GFP was carried out using heterogenous cell populations of interphase cells. To determine if altered IP_8_ levels also affected targeting of kinetochore proteins in M-phase, we analyzed Mal2-GFP kinetochore fluorescence in late anaphase cells and found that in cells without IP_8_ the Mal2-GFP kinetochore signal was also increased significantly (Figure S3B).

### IP_8_, and not IP_7_, controls kinetochore targeting of *S. pombe* CCAN components

Decreasing or eliminating IP_8_ through heterologous expression of the Asp1 pyrophosphatase variant Asp1^365-920^ or via expression of endogenous *asp1*^*D333A*^ (kinase-dead) resulted in an increase of the *S. pombe* CCAN members Mal2 and Fta2 at the kinetochore. However, one important consequence of decreasing IP_8_ levels in this fashion is a concomitant increase in IP_7_ (Figure 3A and 5A). Thus, we needed to determine if the increase in IP_7_ levels might have a role in kinetochore targeting of *S. pombe* CCAN components.

To answer this question, we utilized a genetic strategy. First, we eliminated cellular IP_6_, the fully mono-phosphorylated form of *myo*-inositol that serves as the precursor of yeast IPPs. IP_6_ is generated by Ipk1-mediated phosphorylation at position C2 of the inositol ring (diagram in Figure 6A, top panel). Cells which have the *ipk1*^*+*^ gene deleted, are unable to generate IP_6_, IP_7_ and IP_8_, and instead accumulate large amounts of the Ipk1 substrate IP_5_, along with unconventional IPPs not detected in wild-type *S. pombe* cells (Figure 6A, middle panel) [96]. In an *ipk1Δ* strain, the Mal2-GFP kinetochore signal at the kinetochore was significantly increased (Figure 6B), similar to what we observed for the *asp1*^*D333A*^ strain (Figure 5C). Thus, in both the *ipk1Δ* and the *asp1*^*D333A*^ strains, which have very different IP_7_ levels but share an inability to generate IP_8_, kinetochore targeting of Mal2-GFP was increased.

**Figure 6.**
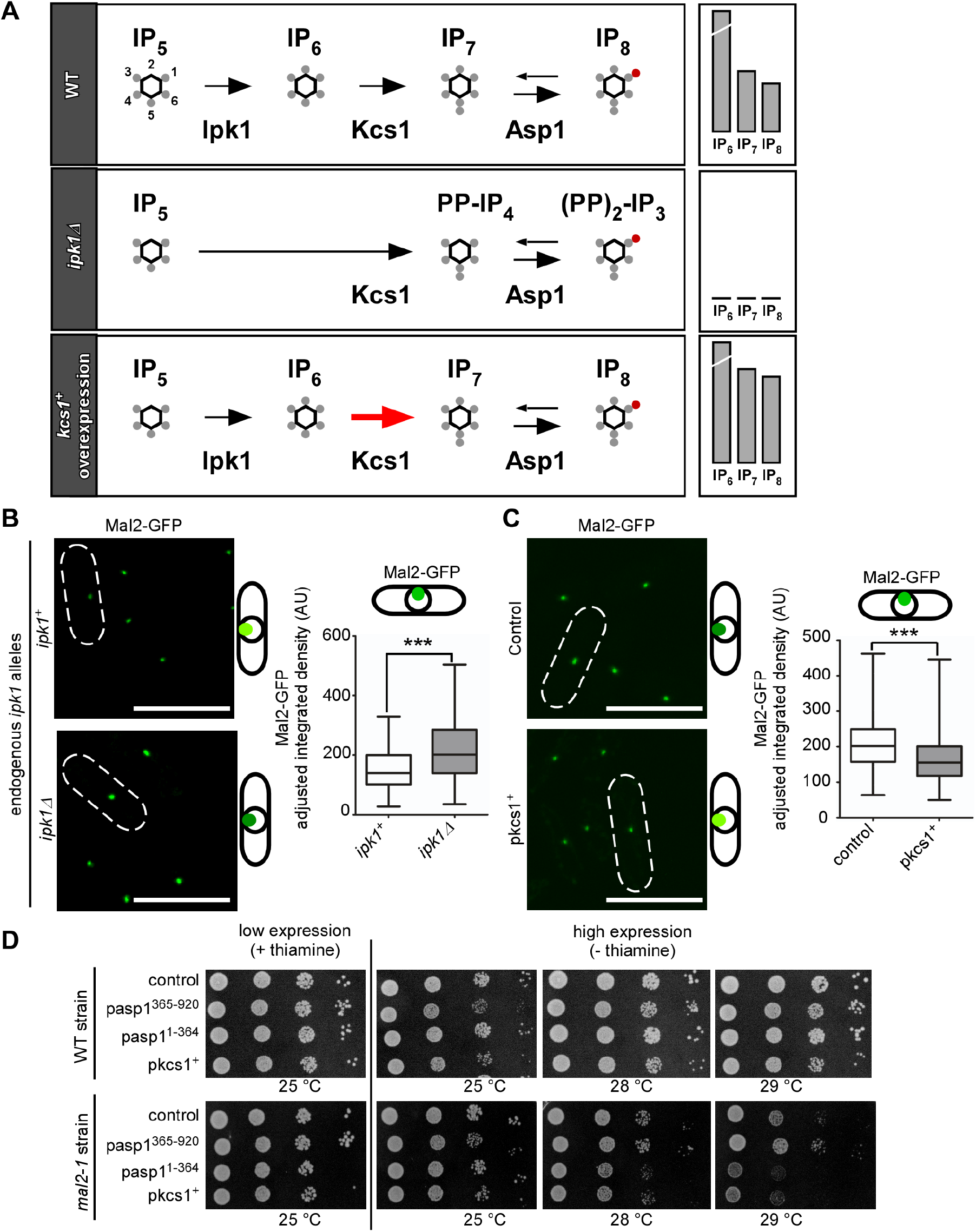
Alterations of the IP_8_ synthesis pathway reveal that kinetochore targeting depends on IP_8_ and not IP_7_ levels. **(A)** Schematic representation of part of the wild-type IP/IPP pathway from IP_5_ to IP_8_, and the changes in IP_7_ and IP_8_ in a *S. pombe ipk1Δ* strain [96] and a wildtype *S. cerevisiae* strain overexpressing *KCS1* (red arrow) based on [99]. **(B)** Left: Live-cell images of *ipk1*^*+*^ and *ipk1Δ* strains endogenously expressing *mal2*^*+*^*-gfp*. Cells were grown at 30 °C. Scale bars, 10 µm. Right: Quantification of Mal2-GFP kinetochore fluorescence signals in each strain. Mean and SD: *ipk1*^*+*^ = 152.3 AU ± 65.86; *ipk1Δ* = 213.1 AU ± 93.54. Number of kinetochore signals counted; *ipk1*^*+*^ = 243; *ipk1Δ* = 183. ***, *p* < 0.0001; Mann-Whitney U-Test. N = 2. **(C)** Left: Live-cell images of a wildtype strain transformed with either control plasmid or a plasmid harboring *kcs1*^*+*^overexpressed via the *nmt41* promoter. Cells were grown at 30°C. Scale bars, 10 µm. Right: Quantification of Mal2-GFP fluorescence signals in each condition. Mean and SD: control plasmid = 205.5 AU ± 65.8; *pkcs1*^*+*^ = 164.6 AU ± 63.39. Number of kinetochore signals counted; control n = 594; pkcs1^+^ n = 609; ***, *p* < 0.0001; Mann-Whitney U-Test. N = 2. **(D)** Serial dilution patch test (10^4^-10^1^ cells) of wildtype and *mal2-1* strains transformed with the indicated plasmids and grown at 25 to 29°C and with/without thiamine for 4-8 days. One of n = 2-3 sets shown.

Next, we targeted the next enzyme in the IPP biosynthetic pathway, Kcs1, which uses IP_6_ as a substrate to generate IP_7_ (Figure 6A, top panel). In contrast to *S. cerevisiae*, where deletion of *KCS1* is viable [97], *S. pombe kcs1*^*+*^ is an essential gene [98], (our analysis). Thus, we perturbed IP_7_ levels by overexpression of *kcs1*^*+*^ on a plasmid via the *nmt41* promoter. Plasmid-borne overexpression of *KCS1* in budding yeast has been shown to increase both IP_7_ and IP_8_ levels [99] (diagrammatically shown in Figure 6A, bottom panel). If increased IP_7_ levels gives rise to higher levels of Mal2 kinetochore targeting, we would expect that overexpression of *kcs1*^*+*^ phenocopies the result observed for the *asp1*^*D333A*^ strain (i.e., no Asp1 kinase activity increases intracellular IP_7_ levels compared to a wild-type strain). Instead, Mal2-GFP kinetochore targeting was decreased in cells overexpressing *kcs1*^*+*^ (Figure 6C), likely due to higher than wild-type IP_8_ levels in these cells. Furthermore, we analyzed the impact of *kcs1*^*+*^ overexpression on the growth of a *mal2-1* strain and found that it resulted in growth reduction at the semi-permissive temperature similar to *mal2-1* cells expressing asp1^1-364^ on a plasmid (Figure 6D). As both types of yeast transformants have increased IP_8_ but opposite levels of IP_7_, we conclude that kinetochore targeting of *S. pombe* CCAN components is solely IP_8_ dependent.

## Discussion

In this study we have demonstrated that Asp1 kinase function has a much broader impact on mitotic processes than previously determined: in a dose-dependent manner IP_8_ is required for entry into mitosis, spindle formation and function [39], and kinetochore architecture. Complete absence of IP_8_ led to defects in chromosome transmission fidelity resulting in aneuploidy and polyploidy [36, 39]. Intriguingly, increasing IP_8_ beyond physiological levels improved transmission of a chromosome above that of a strain with wild-type IP_8_ levels, proposing that IP_8_ is an important player in chromosome transmission fidelity. Using kinetochore targeting of Mal2 as a tool, we have now also demonstrated that it is specifically the IPP IP_8_ that modulates mitosis.

In accordance with the importance of IP_8_ for entry and progression through mitosis, we found that IP_8_ levels are increased at the G2/M boundary but not in G1 or at the metaphase-anaphase A transition. Our analysis does not exclude that IPP levels are cell-cycle regulated in other cell cycle phases apart from G2/M. Indeed, IPP levels change in *S. cerevisiae* cells during progression through S-phase [100]. How such an alteration of IPP levels is regulated in vivo, is unknown. Our finding that only IP_8_ but not IP_7_ levels are elevated at G2/M suggests that (i) it is specifically an IP_8_ increase that is required at this cell cycle stage and (ii) this alteration might be due to downregulation of the Asp1-pyrophosphatase domain and not by other phosphatase activities [101-104].

IP_8_ is required for mitotic entry in a dose-dependent manner. At the permissive temperature for the single mutant *cdc25-22* strain, double mutant *cdc25-22 asp1*^*D333A*^ (no IP_8_) cells showed all phenotypes indicative of a delay at the G2/M transition. The presence of more than the physiological levels of IP_8_ had the opposite effect and partially rescued the *cdc25-22* temperature sensitive growth phenotype. How IP_8_ modulates mitotic entry on a molecular basis will be determined in future studies. However, it has been shown recently that the non-growth phenotype of strains with a deletion of *plo1*^*+*^, which encodes the essential *S. pombe* Polo-like kinase [105] can be rescued by two ways, both of which increase intracellular IP_8_ levels: mutations in the Asp1 pyrophosphatase domain or presence of non-functional Aps1, a member of the DIPP nudix phosphohydrolases that regulate IPP levels [7, 103, 106-108]. As extra IP_8_ is able to bypass the requirement for Polo kinase [106] and as this kinase regulates mitotic entry, in part by activating Cdc25 (reviewed in [109]), we speculate that IP_8_ does not modulate entry into mitosis via Cdc25.

Commitment to mitosis results in the formation of the bipolar spindle and the bioriented attachment of sister chromosomes via their kinetochores to MTs from the opposing spindle poles. We now demonstrate that kinetochore targeting of components of the fission yeast CCAN subcomplex [87] is subject to control by IP_8_ levels. Reduction of IP_8_ in temperature-sensitive CCAN mutant strains *mal2-1, fta2-291* and *mis6-302* partially rescued the non-growth phenotype at higher temperatures, while increased intracellular IP_8_ had the opposite effect. Kinetochore targeting of Mal2-1, wild-type Mal2 and Fta2 in strains with different IP_8_ levels demonstrated an inverse relationship between IP_8_ levels and kinetochore association of these proteins. Furthermore, reduction of IP_8_ increased kinetochore-localized Mal2 beyond that seen in wild-type cells suggesting that cellular pools of Mal2-GFP exceed the amount of proteins localized at the kinetochore; a characteristic also shown for human CENP-Q [58]. What could be the role of IP_8_-mediated kinetochore targeting of fission yeast CCAN components? A characteristic of the CCAN from yeast to humans is that it is present at the centromere during the entire cell cycle and CCAN subunits show a hierarchical set-up (reviewed in [110]). However, abundance of individual components of the CCAN at the kinetochore can vary quite dramatically. For example, the levels of human CCAN members CENP-C/H/T in the kinetochore are 50% increased after nuclear envelope breakdown, while the CCAN member CENP-N exhibited the opposite behavior [111]. Artificial perturbations of IP_8_ levels by using Asp1 variants uncovered the ability of IPPs to modulate kinetochore architecture. However, the finding that IP_8_ levels change at the G2/M transition suggests that this also occurs under physiological conditions. We have assayed just a few members of the *S. pombe* CCAN for IP_8_-dependency but as CCAN components depend on each other for recruitment, it is possible, that changes in the CCAN via IP_8_ are substantial.

Kinetochore targeting of numerous kinetochore proteins is regulated by phosphorylation/dephosphorylation events [112], reviewed in [113]. IPPs can modulate protein function via pyrophosphorylation of a pre-phosphorylated serine residue or via binding to SPX-domain-containing proteins. As only 6 *S. pombe* proteins with an SPX domain exist (https://www.pombase.org/) all with a proven or inferred role in phosphate homeostasis, we propose that kinetochore targeting of *S. pombe* CCAN components is mediated by pyrophosphorylation of a target protein/proteins.

## Supplementary Materials

The following supporting information is available:

Table S1: Strains used in this study; Table S2: List of reagents and plasmids; Supplementary Methods; Figure S1: IP_7_ and IP_8_ levels are similar in wild-type and *cdc25-22* strains grown at 25 °C; Figure S2: Growth phenotype of kinetochore mutant strains expressing *asp1*^*1-364*^ or *asp1*^*365-920*^ from a plasmid via the *nmt1* promoter; Figure S3: Effect of change on IP_8_ levels in Fta2-GFP kinetochore targeting in interphase cells and Mal2-GFP late mitotic cells.

References [114-118] are used only in the Supplementary Materials.

## Author contributions

Conceptualization, Natascha Kuenzel and Ursula Fleig; Formal analysis, Natascha Kuenzel, Abel Alcázar-Román, Adolfo Saiardi and Simon Bartsch; Funding acquisition, Adolfo Saiardi, Dorothea Fiedler and Ursula Fleig; Investigation, Natascha Kuenzel, Abel Alcázar-Román, Adolfo Saiardi, Simon Bartsch and Sarune Daunaraviciute; Methodology, Natascha Kuenzel, Abel Alcázar-Román, Adolfo Saiardi, Simon Bartsch, Sarune Daunaraviciute, Dorothea Fiedler and Ursula Fleig; Project administration, Ursula Fleig; Supervision, Adolfo Saiardi, Dorothea Fiedler and Ursula Fleig; Writing – original draft, Natascha Kuenzel, Abel Alcázar-Román and Ursula Fleig; Writing – review & editing, Natascha Kuenzel, Abel Alcázar-Román and Ursula Fleig.

## Data availability statement

Data is contained within the article or supplementary material.

## Acknowledgements

We thank Boris Topolski and Corinna Braun for help with the images in Figures 3D and S3A. Eva Walla for help with the patch tests shown in Figure 3B, 6D and S2C. Melanie Süssmilch for help with Figures 2F-I (all Heinrich-Heine-University, Düsseldorf). We thank the Center for Advanced Imaging (CAi) at the Heinrich-Heine-University for help with imaging and macro-based image analysis. We furthermore thank Kathleen Gould, Susan R. Wente, Mitsuhiro Yanagida, Yasushi Hiraoka, Takashi Toda, Paul Nurse and the National Bioresource Project (NBRP/YGRC) for *S. pombe* strains and plasmids used in this work. Many thanks to Kathleen Gould for the Asp1 antibody.

