## Supplementary Information for "Inositol pyrophosphate-controlled kinetochore architecture and mitotic entry in *S. pombe*"

### Title

Abel R. Alcázar-Román[a]

Adolfo Saiardi[b]

Simon M. Bartsch[c, d]

Sarune Daunaraviciute[a]

Dorothea Fiedler[c, d]

Ursula Fleig[a,*].

[a] Eukaryotic Microbiology, Institute of Functional Microbial Genomics, Heinrich-Heine-University, Universitätsstrasse, Düsseldorf, 40225, Germany.

[b] Medical Research Council Laboratory for Molecular Cell Biology, University College London, Gower St, London, WC1E 6BT, United Kingdom.

[c] Leibniz Forschungsinstitut für Molekulare Pharmakologie, Robert-Rössle-Straße 10, Berlin 13125, Germany

[d] Humboldt-Universität zu Berlin, Institut für Chemie. Brook-Taylor-Straße 2, 12489 Berlin, Germany

[e] Present address: IUF - Leibniz Research Institute for Environmental Medicine

### Corresponding Author


Supplementary Tables, Methods, Figures and Legends

**Table S1: Strains used in this study.**

| Name and Genotype | Source |
|---|---|
| *cdc25-22 h*$^{+}$ | P. Nurse |
| *cdc10-129 leu1-32 h*$^{+}$ | P. Nurse |
| *cut9-665 leu1 h*$^{-}$ | M. Yanagida |
| *mis6-302 leu1-32 h*$^{-}$ | M. Yanagida |
| MY6105: *mis14-634 leu1 h*$^{-}$ | M. Yanagida |
| KSH060: *ndc80-21::kan*$^{R}$ *leu1 ura4 h-* | Takashi Toda |
| KG425: *his3-D1 ade6-M210 leu1-32 ura4-D18 h*$^{-}$ | K. Gould |
| UFY1065: *mis15-68 leu1-32 ura4-D18 ade6-M210 h*$^{-}$ | Lab Collection (original from M. Yanagida) |
| UFY1057: *mis17-362 his3-D1 leu1-32 ura4-D18 ade6-M210 h*$^{+}$ | |
| UFY1056: *nuf2-1::ura4*$^{+}$ *ura4-D18 leu1-32 ade6-M210 his3-D1, h*$^{-}$ | Lab Collection (original from Y. Hiraoka) |
| UFY518: *fta2-GFP::kan*$^{R}$ *ade6-210 leu1-32 ura4-D6, h*$^{-}$ | Lab Collection |
| UFY547: *mal2-1-GFP::kan*$^{R}$ *ade6-M210 leu1-32 ura4-D18 h*$^{-}$ | Lab Collection |
| UFY852: *mal2-1 leu1-32 ade6-M210 ura4-D18 h*$^{-}$ | Lab Collection |
| UFY1048: *fta2-291::his3*$^{+}$ *his3*$^{-}$*, leu1-32 ura4-D18 ade6-M210 h*$^{-}$ | Lab Collection |
| UFY1058: *mal2*$^{+}$*-GFP::kan*$^{R}$ *ade6-M210 leu1-32 ura4-Dx his3-D1 h*$^{-}$ | Lab Collection |
| UFY1511: *asp1*$^{D333A}$*::kan*$^{R}$ *his3-D1 ade6-M210 leu1-32 ura4-D18 h*$^{+}$ | Lab Collection |
| UFY1577: *mal2-1 ade6-M210 leu1-32 ura4-D18 h*$^{-}$ | Lab Collection |
| UFY1027: *spc7-30/his3*$^{+}$ *his3-D1 ade6-M216 leu1-32 ura4-D18 h*$^{+}$ | Lab Collection |
| UFY2345: *mal2*$^{+}$*-GFP::kan*$^{R}$ *asp1*$^{H397A}$*::kan*$^{R}$ *leu1-32 ura4-D18 his3-D1* | This study |
| UFY2346: *mal2*$^{+}$*-GFP::kan*$^{R}$ *asp1*$^{D333A}$*::kan*$^{R}$ *leu1-32 ura4-18 his3-D1* | This study |
| UFY2349: *asp1*$^{H397A}$*::kan*$^{R}$ *mal2-1-GFP::kan*$^{R}$ *leu1-32 ura4-D18 h*$^{+}$ | This study |
| UFY2351: *asp1*$^{D333A}$*::kan*$^{R}$ *mal2-1-GFP::kan*$^{R}$ *leu1-32 ura4-D18* | This study |
| UFY2370: *fta2-GFP::kan*$^{R}$ *asp1*$^{H397A}$*::kan ade6-M210 leu1-32 ura4-D18* | This study |
| UFY2372: *fta2-GFP::kan*$^{R}$ *asp1*$^{D333A}$*::kan*$^{R}$ *his3-D1 ade6-M210 leu1-32 ura4-D18* | This study |
| UFY3009: *mal2-1-GFP::kan*$^{R}$ *leu1-32 ura4-D18 ade6-M210 h*$^{-}$ | This study |
| UFY3010: *mal2-1-GFP::kan*$^{R}$ *leu1-32 ura4-D18 ade6-M210*$^{+}$ | This study |
| UFY3095: *asp1*$^{D333A}$*::kan*$^{R}$ *cdc25-22 his3-D1? ade6-M210? leu1-32? ura4-D18? h*$^{-}$ | This study |
| UFY3098: *asp1*$^{H3973A}$*::kan*$^{R}$ *cdc25-22 his3-D1? ade6-M210? leu1-32? ura4-D18? h*$^{-}$ | This study |
| UFY3316: *mal2-GFP::kan*$^{R}$ *ipk1Δ::ura4*$^{+}$ *leu1-32 ura4-D18 ade6-M210 h*$^{-}$ | This study (derived from a strain from S. Wente) |

**Table S2. List of reagents and plasmids**

| **Antibodies** | |
|---|---|
| α-Tat1 (IF) | [65] |
| α-mouse Alexa Fluor®488 (IF) | Thermo Fisher Scientific |
| α-GFP (Western Blot) | Roche |
| α-GAPDH (Western Blot) | Sigma Aldrich |
| α-γ-Tubulin (Western Blot) | Sigma Aldrich |
| α-GFP (ChIP) | Molecular Probes |
| α-Asp1 (Western Blot) | Gift from Kathy Gould |
| **Oligonucleotides** | |
| cen1/3 ChIP forward 5’ CAGACAATCGCATGGTACTATC 3’ | Adapted from [71, 72] |
| cen1/3 ChIP reverse 5’ AGGTGAAGCGTAAGTGAGTG 3’ | Adapted from [71, 72] |
| act1 ChIP forward 5’ CCCAAATCCAACCGTGAGAAGATG 3’ | Adapted from [71, 72] |
| act1 ChIP reverse 5’ CCAGAGTCCAAGACGATACCAGTG 3‘ | Adapted from [71, 72] |
| **Plasmids** | |
| pJR2-3XL | [60] |
| pJR1-41XL | [60] |

| | |
|---|---|
| pUF672: pJR2-3XL -*asp1*$^{1\text{-}364aa}$ | [6,60] |
| pUF916: pJR2-3XL-*asp1*$^{365\text{-}920aa}$ | [6,60] |
| pUF148: pREP3XL-*mal2*$^{+}$ | Lab collection [60] |
| pUF503: pJR2-3XL-*fta2*$^{+}$ | Lab collection [60] |
| pUF1035: pJR2-41XL-*mis15*$^{+}$ | Lab collection [60] |
| pUF1489: JR1-41XL-*kcs1*$^{+}$ | This study [60] |
| pUF1027: pREP3XL-NLS-GFP | This study [60] |
| pUF1577 SPX$^{ScVtc2}$GFP | This study [60] |

**Supplementary Methods**

Quantification of fluorescence signals

For manual analysis a MIP picture was created from 15-20 z-stacks. A square of a defined area was drawn around the kinetochore and the mean grey value within the square was measured. The value measured in a square of the same size within the background of the picture was subtracted. For kinetochore signals in mitosis both signals were combined. The MIP image was used for measurement of the signal intensity of GFP fluorescence with ImageJ 1.44 (NIH). For Figure S3B, cells with two clearly separated kinetochore signals (5-10 µm distance) were chosen manually.

For macro-based fluorescence analysis z-stacks (equal within datasets) were recorded. For automated analysis FiJi1.51w was used. A MIP picture was created and ROIs (regions of interest) defined using the following macro for spinning-disk images:

```
run("Duplicate...", " ");
run("8-bit");
run("Subtract Background...", "rolling=50");
run("Maximum...", "radius=2");
run("Threshold...");
waitForUser("thr", "setzit");
setOption("BlackBackground", false);
run("Convert to Mask");
run("Analyze Particles...", "size=0.20-1.00 circularity=0.80-1.00 display exclude clear include
add");
```

For LSM images another macro was used:

```
run("Duplicate...", " ");
run("Subtract Background...", "rolling=5");
setAutoThreshold("Default dark");
//run("Threshold...");
```

```
setThreshold(value1, value2);
//setThreshold(value1, value2);
setOption("BlackBackground", false);
run("Convert to Mask");
run("Analyze Particles...", "size=0.05-Infinity circularity=0.20-1.00 display exclude clear
include add");
```

The threshold (window that opens in spinning disk macro or value1 and value2 in LSM macro) was set for each experiment so that most GFP signals were defined as a ROI while as little artifacts as possible were marked. The same thresholds were used for a given experiment set. The signal intensity was measured by transferring the saved ROI-set to the unedited MIP and using the measure option. The value used for quantification was the Integrated Density (IntDen) value as it accounts for the area and the mean gray value of the ROI. For the adjusted integrated density value, the background signal in an area equal to that of the ROI was subtracted from each value. The number of quantified signals is given as number of kinetochore signals in the figure legends.

Quantification and Statistical Analysis

Statistics

The number of replicates and statistical tests used are shown in the figure legends. Statistics were performed with the GraphPad Prism software. Significant outliers were removed using Grubbs' test.

Quantification of qChIP cen1/3 over actin values

The calculation of qChIP data was performed as follows [92]:

1) ΔCt values for the cen1/3 and act qPCRs were determined:

$\Delta Ct = Ct_{IP} – (Ct_{input} – \log(10;2))$

2) % input values for the cen1/3 and act qPCRs were calculated:

% input = $(2^{-\Delta Ct})$

3) % enrichment values for cen1/3 over act:

enrichment values cen1/3 over act = (% input cen1/3)/(% input act)

#### Supplementary Figures

**Figure S1. $IP_7$ and $IP_8$ levels are similar in wild-type and *cdc25-22* strains grown at 25 °C. (A)** Typical HPLC profiles of soluble inositol phosphates and pyrophosphates extracted from wild-type and *cdc25-22* strains grown at 25 °C (not arrested). Magnification of the $IP_7$

and $IP_8$ peaks is shown on the right-hand side. **(B)** Quantification of $IP_7$ levels relative to $IP_6$. Wildtype = 9.75 ± 1.626; *cdc25-22* = 9.3 ± 2.546 **(C)** Quantification of $IP_8$ levels relative to $IP_6$. Wildtype = 12.15 ± 2.051; *cdc25-22*: 10.65 ± 0.495. For (B) and (C), n.s. = not significant. n = 2.

**Figure S2. Growth phenotype of kinetochore mutant strains expressing *asp1$^{1\text{-}364}$* or *asp1$^{365\text{-}920}$* from a plasmid via the *nmt1* promoter**. **(A)** Result of patch test analysis of the indicated transformants grown at 25 °C and at non-permissive temperatures [114-118]. The phenotype was scored under high expression of the Asp1 variants. -: growth as for the transformants with a control plasmid; arrow down: negative effect on growth. Size of the arrow increases with severity of phenotype. Arrow up: non-growth phenotype at non-permissive temperature is rescued. **(B)** Serial dilution patch test of *mis6-302* and *mis15-68* strains transformed with the indicated plasmids and grown at the indicated temperatures for 4-8 days depending on the incubation temperature. Pasp1$^{1\text{-}364}$ and pasp1$^{365\text{-}920}$, plasmids with *nmt1* driven expression of the Asp1 kinase or pyrophosphatase, respectively. Growth on media with thiamine led to low expression of these Asp1 variants, while growth on media without thiamine, resulted in high expression of the Asp1 variants. One of n = 1-3 sets shown. **(C)** Serial dilution patch test experiments of *mal2-1-gfp* strains transformed with the indicated plasmids and grown at the indicated temperature for 5 days. pmal2$^{+}$, plasmid with wild-type *mal2$^{+}$* ORF expressed via the *nmt1* promoter. pasp1$^{1\text{-}364}$ and pasp1$^{365\text{-}920}$, plasmids with *nmt1* driven expression of the Asp1 kinase or pyrophosphatase, respectively. *nmt1* transcribed genes show low expression in the presence of thiamine and high expression when no thiamine is present.

**Figure S3. Effect of change on $IP_8$ levels in Fta2-GFP kinetochore targeting in interphase cells and Mal2-GFP late mitotic cells. (A)** Left: Live cell images of the indicated *asp1*-variant strains endogenously expressing *fta2$^{+}$-gfp* and grown at 25˚C. Scale bars, 10 µm. Right: Quantification of Fta2-GFP fluorescence signals: Mean and SD: *asp1$^{+}$* = 243.7 AU ± 107.4; *asp1$^{H397A}$* = 172.5 AU ± 67.43; *asp1$^{D333}$* = 611.9 AU ± 202.1. Number of kinetochore signals counted, *asp1$^{+}$* = 243; *asp1$^{H397A}$* = 281; *asp1$^{D333A}$* = 145; ***, $p < 0.0001$; Mann-Whitney U-Test. **(B)** Left: Live cell images of the indicated *asp1*-variant cells with fully separated chromosomes endogenously expressing *mal2$^{+}$-gfp* grown at 25˚C. Scale bars, 10 µm. Right: Manual quantification of separated Mal2-GFP fluorescence signals in pictures as shown in (A). Mean and SD: *asp1$^{+}$* = 9 AU ± 3.74; *asp1$^{H397A}$* = 9.29 AU ± 5.09; *asp1$^{D333A}$* = 14.34 AU ± 7.3. Number of kinetochore signals counted: *asp1$^{+}$* n = 20; *asp1$^{H397A}$* and *asp1$^{D333A}$* n = 21; **, $p = 0.0058$; t-test with Welch's correction. Average of n = 3 shown.

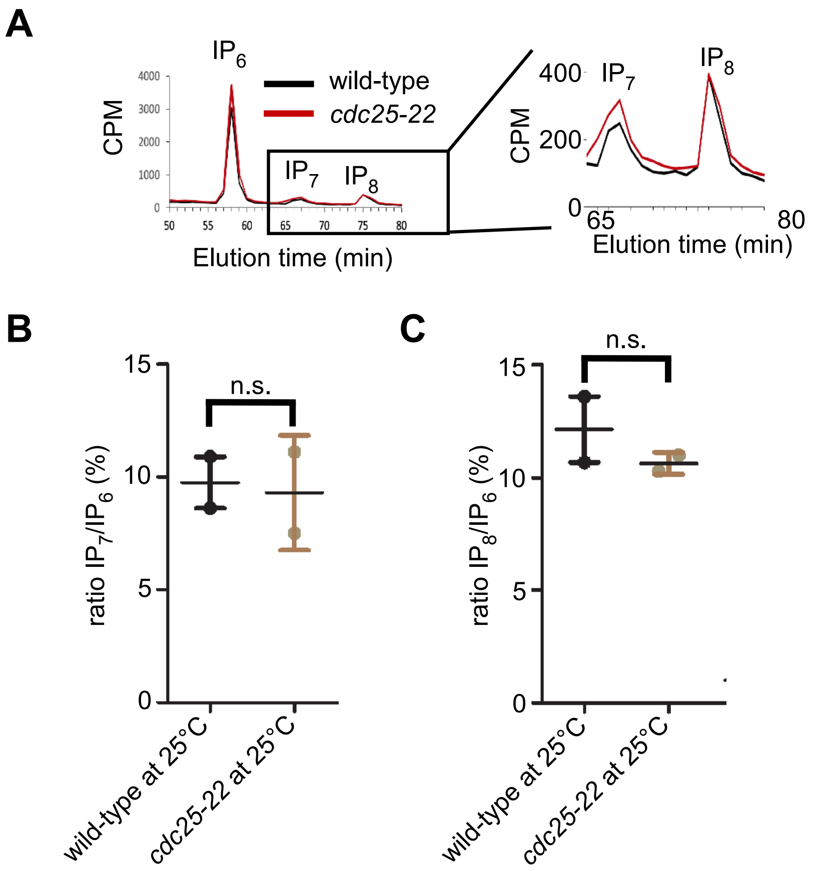
A
IP6
wild-type
cdc25-22
CPM
4000
3000
2000
1000
0
IP7
IP8
50
55
60
65
70
75
80
Elution time (min)
400
200
0
65
80
B
15
10
5
0
n.s.
ratio IP7/IP6 (%)
wild-type at 25°C
cdc25-22 at 25°C
C
n.s.
ratio IP8/IP6 (%)

**Figure S1**

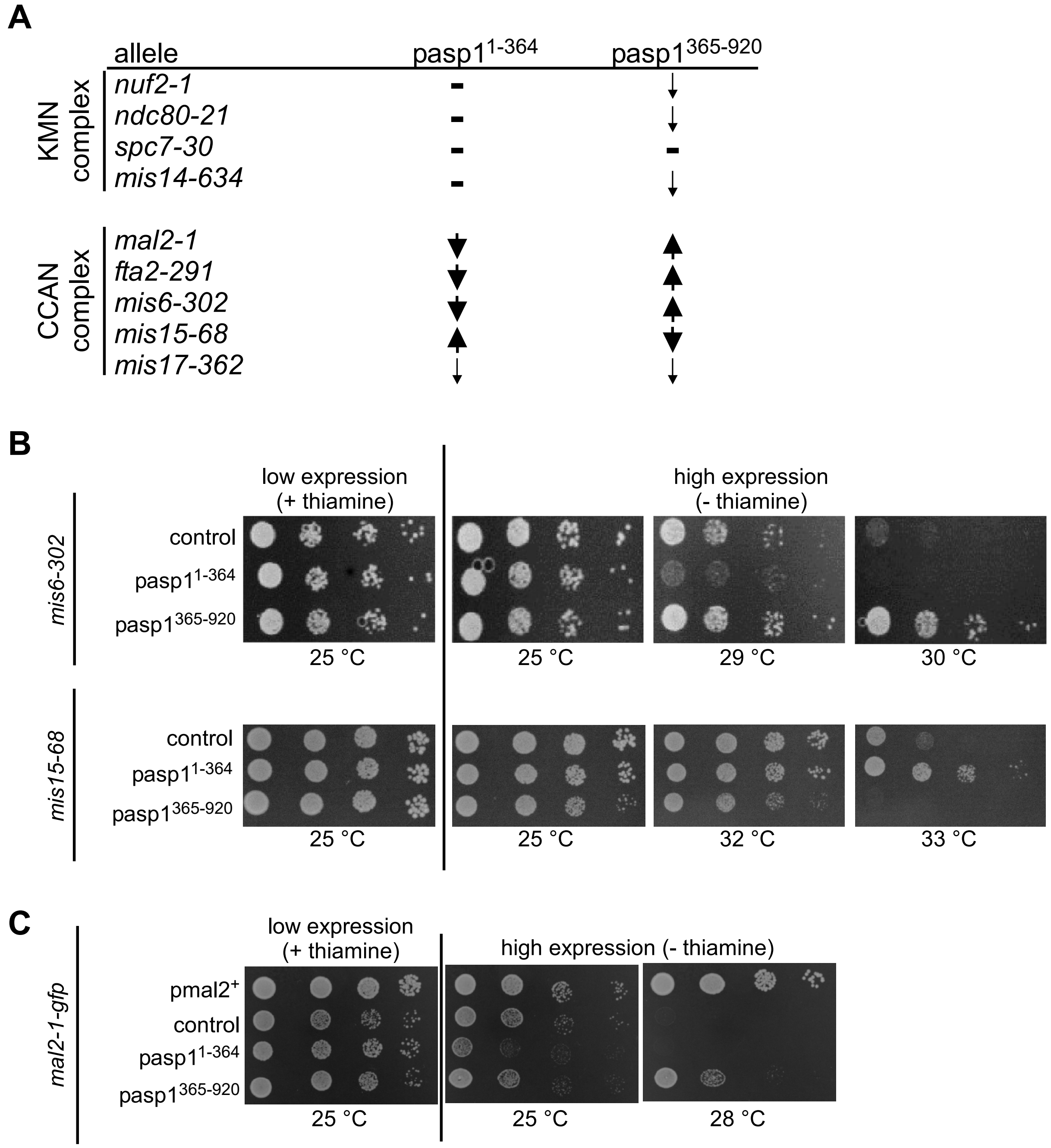
A
allele
pasp1 1-364
pasp1 365-920
KMN complex
nuf2-1
ndc80-21
spc7-30
mis14-634
CCAN complex
mal2-1
fta2-291
mis6-302
mis15-68
mis17-362
B
low expression (+ thiamine)
high expression (- thiamine)
mis6-302
control
pasp1 1-364
pasp1 365-920
25 °C
25 °C
29 °C
30 °C
mis15-68
control
pasp1 1-364
pasp1 365-920
25 °C
25 °C
32 °C
33 °C
C
low expression (+ thiamine)
high expression (- thiamine)
mal2-1-gfp
pmal2+
control
pasp1 1-364
pasp1 365-920
25 °C
25 °C
28 °C

**Figure S2**

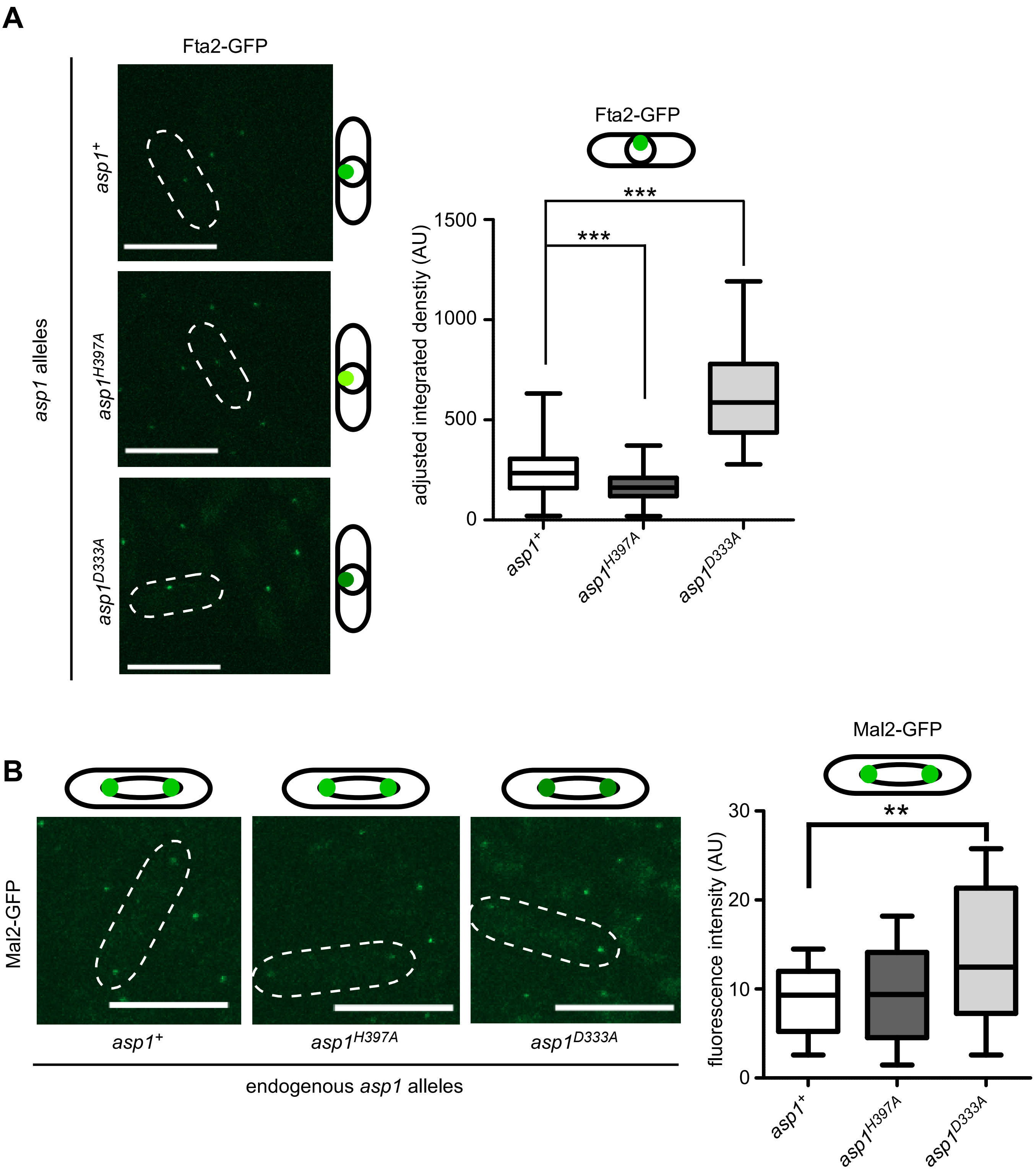
A
Fta2-GFP
asp1 alleles
$asp1^+$
$asp1^{H397A}$
$asp1^{D333A}$
Fta2-GFP
***
***
adjusted integrated densitiy (AU)
1500
1000
500
0
B
Mal2-GFP
Mal2-GFP
**
fluorescence intensity (AU)
30
20
10
0
endogenous asp1 alleles

**Figure S3**
